# CMT3 and SUVH4/KYP silence the exonic retroelement Evelknievel to allow for reconstitution of *CMT1* mRNA

**DOI:** 10.1101/332239

**Authors:** Narendra Singh Yadav, Janardan Khadka, Assaf Zemach, Gideon Grafi

## Abstract

Highlights:

- Silencing of the intragenic Evelknievel (EK) retroelement is maintained by CMT3-KYP/SUVH4 independently of DDM1 and the RdDM pathways.
- Methylation and silencing of EK is required for transcription through the EK retroelement.
- Silencing of EK allows for splicing out of the entire EK and reconstitution of an intact CMT1 mRNA.

**Background:** *CHROMOMEHYLASE1* (*CMT1*) has long been considered a non-essential gene because, in certain *Arabidopsis* ecotypes, the *CMT1* gene is disrupted by the retroelement Evelknievel (EK), inserted within exon 13, or contains frame-shift mutations resulting in a truncated, non-functional protein. Here, we wanted to explore the regulatory pathway responsible for EK silencing in the Ler ecotype and its effect on CMT1 transcription.

**Results:** Methylome databases confirmed that EK retroelement is heavily methylated but methylation is extended toward *CMT1* downstream region. Strong transcriptional activation of EK accompanied by significant reduction in non-CG methylation was found in *cmt3* and *kyp2*, but not in *ddm1* or RdDM mutants. EK activation in *cmt3* and *kyp2* did not interfere with upstream *CMT1* expression but abolish transcription through the EK. We identified, in wild type Ler, three spliced variants in which the entire EK is spliced out; one variant (25% of splicing incidents) facilitates proper reconstitution of an intact *CMT1* mRNA. We could recover very low amount of the full length *CMT1* mRNA from WT Ler and Col but not from *cmt3* mutant..

**Conclusions:** Our findings highlight CMT3-SUVH4/KYP as the major pathway silencing the intragenic EK *via* inducing non-CG methylation. Furthermore, retroelement insertion within exons (e.g., *CMT1*) may not lead to a complete abolishment of the gene product when the element is kept silent. Rather the element can be spliced out to bring about reconstruction of a very low level of an intact, functional mRNA and possibly to retrieval of an active protein.

## Background

Because of their abundance and the potential to modify/mutate the genome [1,2], transposable elements (TEs) are subjected to multiple layers of epigenetic control to ensure their silencing. Accordingly, silencing of TEs is accomplished by complementary epigenetic mechanisms that include chromatin compaction associated with DNA methylation and histone modification often mediated by small RNAs [3]. Furthermore, recent study revealed an interesting correlation betweem TE location: near a gene, within a gene, in a pericentromere/TE island, or at the centromere core and the regulatory mechanism underlying its silencing (Sigman and Slotkin, 2016). In *Arabidopsis*, methylation in the context of CpG is maintained by METHYLTRANSFERASE 1 (MET1), a homolog of the mammalian Dnmt1 [4,5]; in *met1* mutant, most CpG methylation is lost [6,7]. Methylation in the CHG (H represent A, C or T) context is maintained by a plant-specific DNA methyltransferase CHROMOMETHYLASE 3 (CMT3) [8,9], which often requires methylation of histone H3 at lysine 9 by H3K9 methyltransferases SUVH4/KYP, SUVH5 and SUVH6 [10]. Genome wide profiling of DNA methylation revealed that CMT3 preferentially methylate transposons including those that are present as single copies within the genome [11]. Notably, single mutants of *met1* or *cmt3* displayed significant accumulation of CACTA transcript [12], the most abundant class II superfamily of plant transposons [13]. However, mobilization of these elements was very limited in single *met1* or *cmt3* mutants, while in *met1 cmt3* double mutant high frequency transposition of these elements was observed [12]. In a recent work, Khan *et al*. [14] showed that the class II transposon Tag1 is essentially silenced by CMT3 *via* gene body CHG methylation; Tag1 was strongly and slightly activated in *cmt3* and *ddm1* (*decrease in DNA methylation 1*), respectively. While maintenance of CG, CHG, and CHH methylation are maintained by MET1, CMT3 and CMT2 [8,9,15], *de novo* methylation in all sequence contexts is essentially established by DOMAINS REARRANGED METHYLTRANSFERASEs (DRM1/DRM2) [16,17]. DRM1/2 mediate *de novo* methylation *via* 23–24-nt small interfering RNAs (siRNAs)-directed DNA methylation (RdDM) pathway. RdDM is a complex mechanism, which involves multiple factors and steps including the formation of double stranded RNA from Pol IV-derived transcripts by RNA-DEPENDENT RNA POLYMERASE 2 (RDR2) and its processing by DICER-LIKE 3 (DCL3) into 23-24-nt siRNAs, which are exported to the cytoplasm [18]. Besides methyltransferases, the chromatin remodeling factor DDM1 appears to play a major role in maintaining cytosine methylation in CpG and non-CG contexts and silencing of genes and transposable elements. Accordingly, mutation of DDM1 has led to significant reduction in global cytosine methylation [19,20] particularly at heterochromatic H3K9me2-enriched regions [21], to activation of certain genes [22] and to reactivation and transposition of retroelements [23,24]. DDM1 appears to provide DNA methyltransferases such as CMT2 access to H1-containing heterochromatin to maintain silencing of TEs in cooperation with the RdDM pathway (25).

*CHROMOMEHYLASE 1* (*CMT1*) – a paralog of *CMT3* - has long been considered a non-essential gene because, in certain *Arabidopsis* ecotypes (e.g., Ler), the *CMT1* gene is disrupted by a single copy retroelement Evelknievel (EK), inserted within exon 13, or contains frame-shift mutations resulting in a truncated, non-functional protein [26]. The EK retroelement contains perfect long-terminal repeats (LTRs), and encodes for a protein containing 1451 amino acids [26]. Here, we wanted to explore the regulatory pathway responsible for EK silencing in the Ler ecotype and how EK silencing affect CMT1 transcription. We report here that CMT3-SUVH4/KYP is the major pathway controlling silencing of the exonic EK *via* inducing non-CG methylation independently of DDM1 and RdDM and that EK silencing is required for reconstitution of intact *CMT1* mRNA.

## Methods

### Plant materials

All *Arabidopsis* lines, wild type Col and Ler, as well as mutants in the Ler background, namely, *ddm1* (Ler background CSHL-GT24941), *cmt3-7* (CS6365, provided by D. Autran) and *kyp2* (CS6367, provided by D. Autran), *rdr2* (provided by Bin Yu), *ago4* (provided by Caroline Dean) and *hen1* (provided by S. Mlotshwa, V. Vance lab) were grown in a controlled growth room under long day photoperiod (16 h light and 8 h dark, light intensity 200 μmoles photons m^-2^ s^-1^) at 22°C ±2 and 70% humidity.

### DNA and RNA isolation, cDNA production and expression analysis

DNA was extracted from plant tissues using Genomic DNA Mini kit (Geneaid, Taiwan). RNA was prepared from leaves, protoplasts or flowers using RNeasy Plant Mini Kit (Qiagen). 1 μg of total RNA was used for cDNA production using the Verso cDNA kit (Thermo scientific) according to the manufacturer’s protocol. The resulting cDNA was subjected to PCR to amplify the Evelknievel using primer set EK-RTF/EK-RTR, AtCOPIA18A using primers 18A-F/ 18A-R, Solo LTR using primer set Solo-F/Solo-R and AtMu1 using primer set AtMu1-F/AtMu1-R. Upstream analysis of *CMT1* expression was performed using two primer sets, ex10-F(P1)/ex11-R(P2) and ex11F(P3)/ex13R(P4) and for down stream CMT1 expression we used ex14-F(P5)/ex16-R(P6). Finally for the analysis of splicing out of the entire EK we used ex13-F(P7)/ex16-R(P6) primer set. Actin or Ubiquitin10 were used as controls (primer set ACT-F/ACT-R and UBQ10-F/ UBQ10-R, respectively) (for primer sequences see Supporting information, Table S1). PCR conditions were 95°C, 2 min; 30-40 cycles of 95°C, 30 s; 60°C, 30 s; 72°C, 30 s; followed by 72°C, 5 min. PCR products were resolved on 1.5% agarose gel stained with ethidium bromide.

### Bisulfite sequencing

Bisulfite conversion was carried out using the Qiagen EpiTech bisulfite kit according to the manufacturer’s instructions on genomic DNA extracted from rosette leaves of WT Ler and various mutant lines in the Ler background using Genomic DNA Mini kit (Geneaid, Taiwan). The reactions with a C to T conversion rate higher than 98% (as determined by the sequencing of 10 clones of an unmethylated control DNA, a 1562 bp PCR product) were used for further analyses. The bisulfite treated DNA was used for PCR amplification of selected target regions of Evelkenivel retroelement using primer sets bsEK5LTR-F1/bsEx14-R1 and bsEKcr-F1/bsEKcr-R1 to amplify promoter and internal EK regions, respectively. To confirm identity of amplified EK fragments, the initial PCR was followed by nested PCR using primer sets bsEK5LTR-F2/bsEx14-R2 and bsEKcr-F2/bsEKcr-R2 to amplify promoter and internal EK regions, respectively. The conditions for both PCR reactions were 95 °C, 2 min; 30 cycles of 95 °C, 30 s, 60 °C, 30 s and 72 °C, 30 s; followed by 72 °C, 5 min. The amplified sequences were separated by agarose gel electrophoresis before purification using the Qiaquick PCR Purification Kit (Qiagen). The purified fragments were ligated into a pJET1.2 cloning vector using a CloneJET PCR Cloning Kit (Thermo Fisher Scientific) and transformed into competent *E. coli* (TOP10 cells). Positive clones were selected by colony PCR followed by plasmid isolation with the PrestoTM Mini Plasmid kit (Geneaid). At least 10 individual clones for each region of each genotype were sequenced using pJET1.2 primers at Macrogen Europe (Amsterdam, The Netherlands). The sequences were analyzed with Kismeth software (27) to obtain the percentage of methylated sites for each sequence context.

### Analysis of DNA methylation by methylation sensitive enzymes

Genomic DNA was digested with methylation sensitive enzymes including *Hpa*II, *Msp*I, *Sau3*AI, *and Bgl*II and subjected to PCR to amplify various EK DNA sequences. The following primers were used (see Supporting information, Table S1): EK-RTF/EK-RTR primer set for recovery after *Hpa*II and *Msp*I digestion, EKSau3-F/EKSau3-R and EKBgl-F/EKBgl-R primer sets for recovery after *Sau*3AI and *Bgl*II digestion, respectively. As a control for chop PCR we used EKcont-F/EKcont-R primer set to amplify EK sequence lacking the abovementioned restriction sites. PCR conditions were 95°C, 5 min; 30-35 cycles of 95°C, 30 s, 60°C, 30 s and 72°C, 30 s; followed by 72°C, 5 min. PCR products were resolved on 1.5% agarose gel stained with ethidium bromide.

### Analysis of CMT1 full length mRNA

RNA was prepared from flowers using RNeasy Plant Mini Kit (Qiagen). 1 μg of total RNA was used for cDNA production using oligo dT primer and the Verso cDNA kit (Thermo scientific) according to the manufacturer’s protocol. One fifth of the resulting cDNA was subjected to first PCR (46 cycles) using Go-Taq polymerase (Promega) to amplify the *CMT1* full length cDNA with CMT1-F and CMT1-R1 primers (Table 1). PCR conditions were 95 °C, 3 min; 46 cycles of 95 °C, 45 s; 55 °C, 1 min; 72 °C, 3 min; followed by 72 °C, 10 min. PCR products were resolved on 1.0% agarose gel stained with ethidium bromide. Thereafter, the nested PCR was performed using CMT1-F and CMT1-R2 primers (Table 1) using 5μl of the first PCR product as template with same first PCR thermal cycler programme and resolved as described above.

## Results

### Evelknievel (EK) is heavily methylated in the Ler genome

We investigated the regulation of the intragene class I, copia-like retroelement Evelknievel (EK), which is inserted in exon 13 of the *CMT1* gene in the Ler but not in the Col genome. We first Screened available methylome database of WT Ler [28] for the methylation pattern of the EK retroelement and found (Figure 1A) that EK is heavily methylated in all cytosine contexts in WT Ler, which is consistent with the data obtained using methylation sensitive enzymes by Henikoff and Comai [26]. In addition, cytosine methylation is extended downstream from EK insertion site into the 3’ end of the *CMT1* gene.

**Fig. 1.**
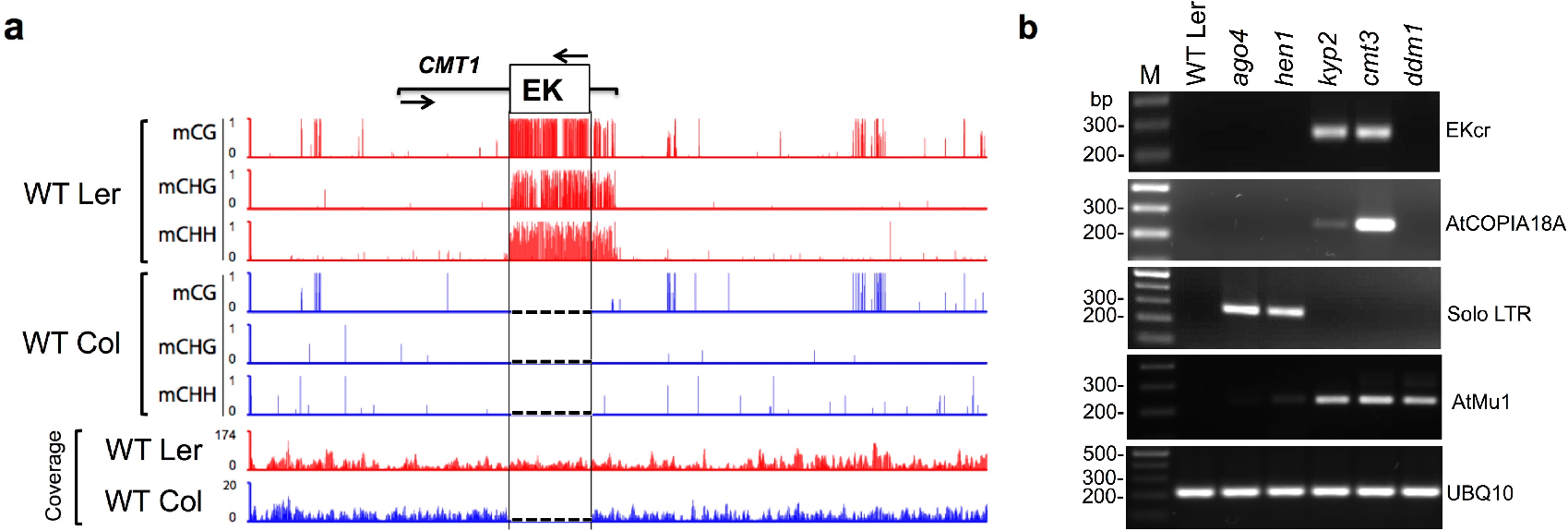
The intragene Evelknievel retroelement is heavily methylated in the Ler genome. **a** The methylation pattern at the indicated cytosine contexts and their position along the EK-CMT1 sequence in WT Ler (red) and WT Col (blue) is shown (GSE34658, Shen *et al*, 2012). Arrows indicate the transcriptional direction of *CMT1* and EK. Note that DNA methylation at all cytosine contexts is extended to *CMT1* 3’ end down stream from the EK insertion site and that EK is absent from WT Col (no coverage) and *CMT1* gene is essentially unmethylated. Broken lines in Col indicate the absence of EK. **b** EK is activated in *cmt3* and *kyp2* but not in *ddm1* or RdDM mutants. Transcriptional activation of EK and other TEs in WT Ler and in various epigenetic mutants. cDNAs were prepared from RNA extracted from leaves of WT Ler, *ago4*, *hen1*, *kyp2*, *cmt3* and *ddm1* and subjected to PCR to amplify EKcr coding region, AtCOPIA18A coding region, Solo LTR and AtMu1. UBQ10 was used as a reference. M indicates DNA molecular size markers.

### EK is expressed in *cmt3* and *kyp2* but not in *ddm1* or RdDM mutants

To gain insight into the regulation of the exonic EK and its effect on *CMT1* transcription we screened several epigenetic mutants in the Ler background for EK activation. We selected mutants affecting DNA and histone methylation, including *ago4* and *hen1* involved in RdDM as well as *ddm1, cmt3 and kyp2.* We generated cDNA from total RNA prepared from leaves of WT Ler, *ago4*, *hen1, cmt3*, *kyp2* and *ddm1* followed by PCR to amplify EK coding region (EKcr). The results showed (Figure 1B) that EK was strongly activated in *cmt3* and *kyp2* mutants but not in WT Ler, *ddm1*, *ago4* and *hen1*. Interestingly, similarly to EK, transcription of AtCOPIA18A reroelements, in which a single copy exist in the Ler genome on chromosome 5, is up-regulated in *kyp2* and *cmt3* mutants but not in WT Ler, *ddm1*, *ago4* and *hen1* (Figure 1B). These results suggest that transcriptional silencing of the retroelements, EK and AtCOPIA18A, is maintained by CMT3 and SUVH4/KYP independently of DDM1 and the RdDM pathway. To further confirm the specificity of TE regulation by SUVH4/KYP-CMT3 pathway, we analyzed the expression of *solo LTR* previously reported to be a target of the RdDM pathway [29] and AtMu1 reported to be activated in *ddm1* mutant [30]. Consistent with previous reports *solo LTR* was restrictively expressed in *ago4* and *hen1* RdDM mutants, while *AtMu1* showed strong expression in *ddm1* as well as in *cmt3* and *kyp2* mutants (Figure 1B).

We analyzed cytosine methylation in various lines by bisulfite sequencing. To this end, genomic DNAs prepared from WT Ler, *hen1*, *ago4*, *rdr2*, *ddm1*, *cmt3* and *kyp2* were treated with sodium bisulfite and the resulting DNAs were used as templates for PCR amplification of EK-5’LTR-CMT1 region and EK coding region (EKcr). PCR fragments were cloned into pJET1.2 and multiple clones from each line were sequenced (Supporting information, BS seq). Cytosine methylation was significantly reduced, particularly in CHG and CHH contexts, in *cmt3* and *kyp2* mutants both in EK-5’LTR-CMT1 and EKcr (Figure 2A and 2B); cytosine methylation, in all contexts, in *ddm1*, *rdr2*, *hen1* and *ago4* mutants were essentially similar to that of WT Ler. Notably, *cmt3* mutant also displayed a significant effect on CG methylation, whereby 60% reduction in methylation was observed; such reduction is not seen in *kyp2* mutant. Similar results were reported previously for *cmt3* mutant displaying also a significant reduction in CG methylation [8]. This is probably due to dependency of CG methylation on non-CG methylated sites, which was observed in several mutants including *kyp suvh5 suvh6* triple mutant and *cmt3* [21]. More prominent effect was observed in the EK-5’LTR-CMT1 where methylation at all cytosine contexts was completely erased in *cmt3* and *kyp2* mutants both in EK 5’LTR and in CMT1 downstream sequence. We further confirmed the EKcr methylation pattern by chop PCR using various methylation sensitive restriction enzymes (Supporting information, Figure S1).

**Fig. 2.**
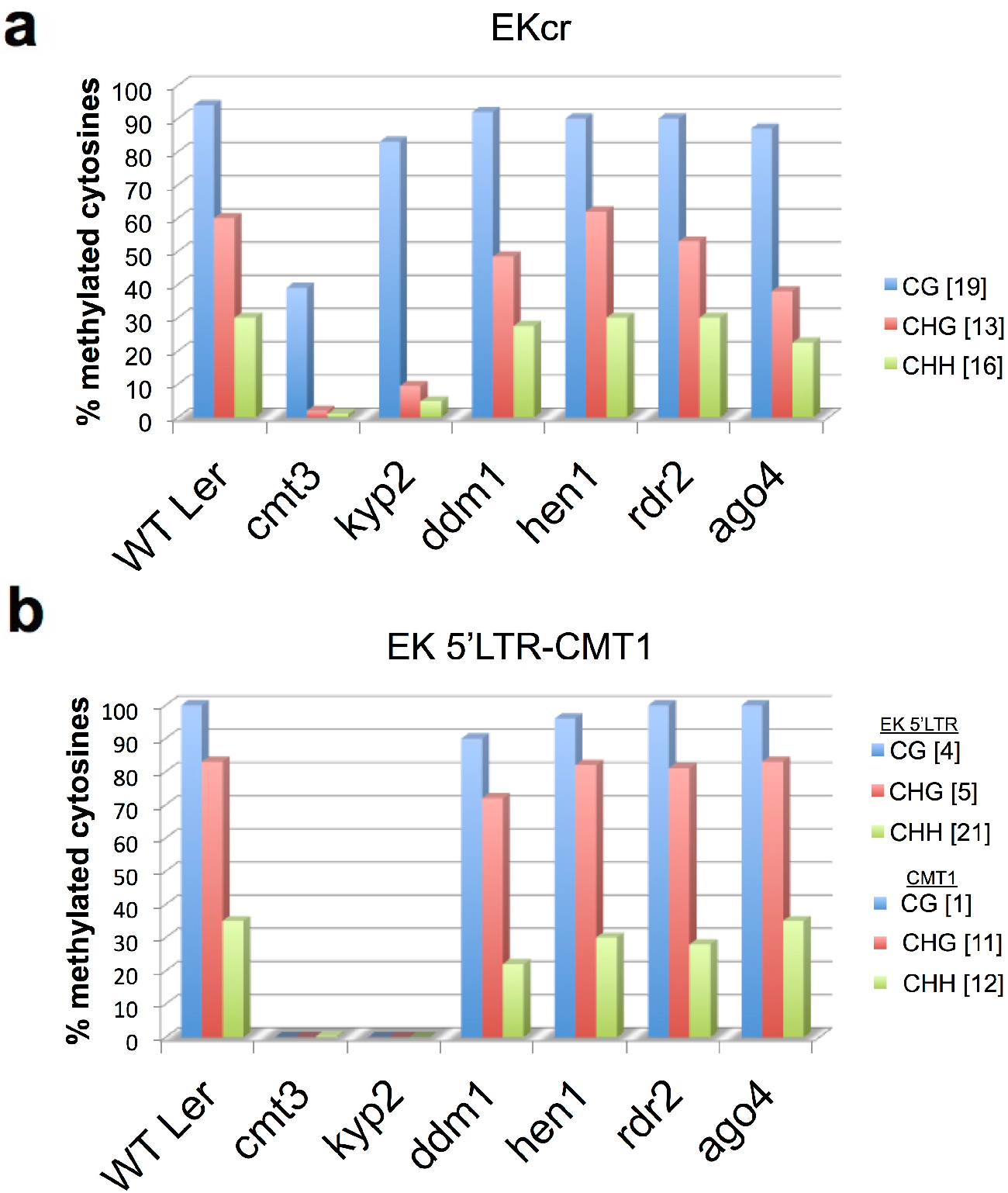
EK Non-CG methylation is significantly and specifically reduced in *kyp2* and *cmt3* mutants. Bisulfite sequencing was performed on genomic DNA prepared from the indicated lines followed by PCR amplification of EKcr (**a**) or the EK 5’LTR-CMT1 (**b**) sequence. Amplified DNA fragments were sub-cloned into pJET1.2 and multiple clones from each line were sequenced (Supporting information, BS-seq) and the percentage of cytosine methylation was determined. The total number of CG (blue column), CHG (brown column) and CHH (green column) sites in amplified DNA sequences is indicated in square brackets on the right.

### Silencing of EK is required for reconstitution of an intact *CMT1* mRNA

We examined how hypomethylation and transcriptional activation of EK affect transcription of the *CMT1* gene. Previously it has been reported that in the *Arabidopsis* No-0 ecotype EK insertion within exon 13 of the *CMT1* gene, had no effect on transcription and splicing upstream the insertion site and the full length cDNA corresponding to the expected size of *CMT1* mRNA could be amplified; although the entire EK is spliced out, the reading frame is shifted resulting in truncated, non-functional protein [26]. To examine *CMT1* expression upstream of the EK insertion site we first generated primers corresponding to exon 10 and exon 11 (Figure 3A, primer set P1/P2) shown previously to amplify CMT1 mRNA fragment from No-0 ecotype [26]. We used cDNA generated from RNA prepared from flowers (where CMT1 is highly expressed [26]) of WT Ler and of various mutant lines; *CMT1* expression in Col flowers was used as a reference. Results showed (Figure 3B) the amplification of a single fragment from WT Ler and mutant lines, which was comparable to that of the Col ecotype. Thus, the upstream *CMT1* gene is transcribed in Ler and mutants similarly to its expression in Col and independently of EK methylation and transcription. Further examination of the upstream *CMT1* transcription using primers corresponding to exon 11 and exon 13 (primer set P3/P4, Figure 3A) revealed the amplification of a single fragment in Col and two fragments, with similar intensities, in WT Ler and mutant lines; one fragment was comparable to that of Col (SP1) and the second was slightly larger (SP2), which appears to be due to intron 12 retention (Figure 3B). We verified the identity of the amplified PCR fragments by sequencing showing that indeed SP2 fragment is a product derived from alternative splicing and is composed of intron 12 (Supporting information, Figure S2). This spliced variant is predicted to have a stop codon within intron 12 that might result in a truncated protein lacking the downstream methylase catalytic domains.

**Fig. 3.**
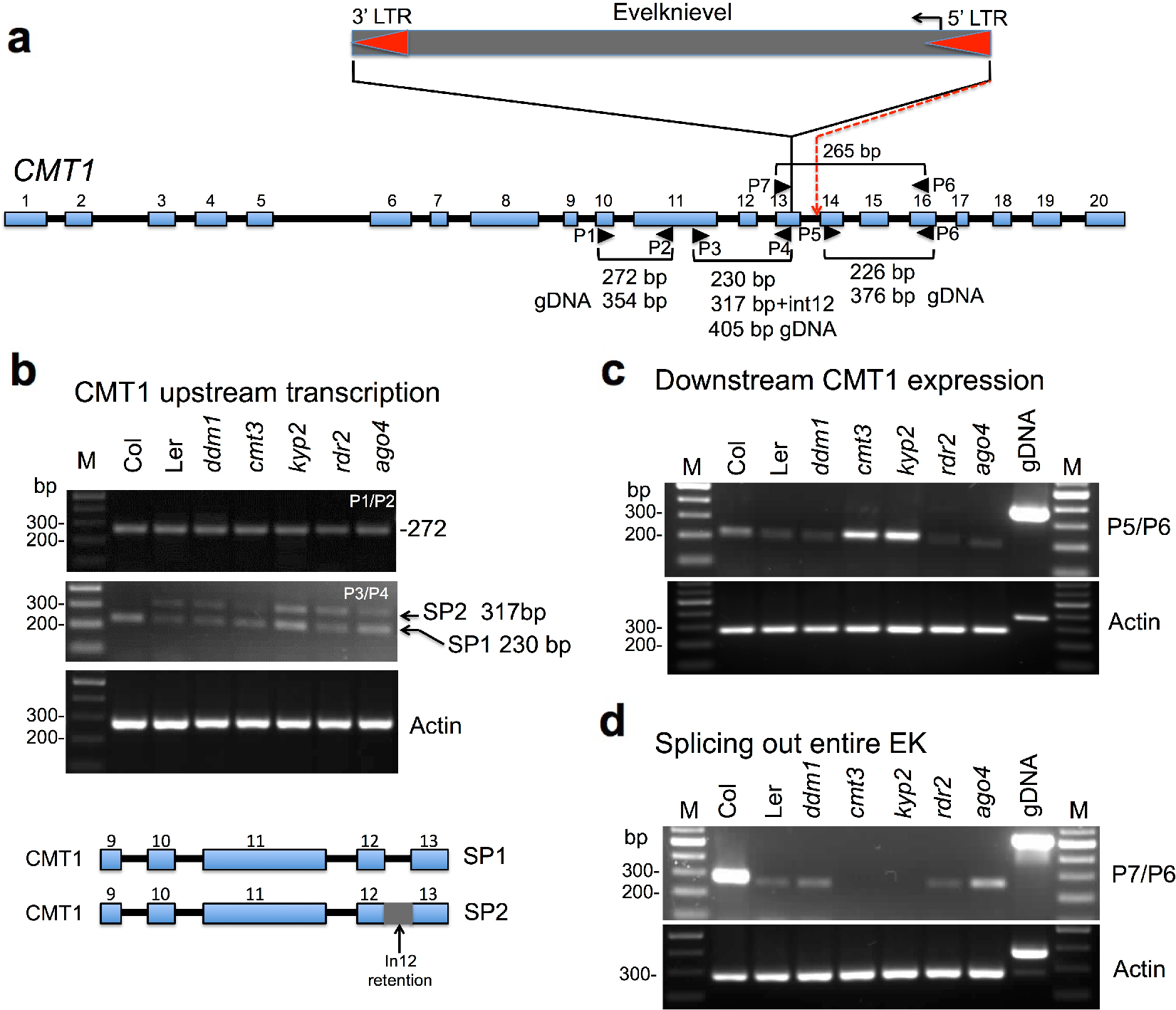
EK transcriptional activation affects *CMT1* downstream expression. **a** Schematic representation of the *CMT1* gene and the insertion of EK within exon 13. *CMT1* exons are numbered and shown as blue boxes and introns as black lines. Primers used to amplify upstream and downstream CMT1 sequences from cDNA are marked by arrowheads and numbered (P1–P7). The red broken line indicates the splice acceptor site of exon 14 (based on 26). EK LTRs are marked by red arrowheads. **b** *CMT1* upstream transcript is alternatively spliced showing intron 12 retention. cDNA were prepared from RNA extracted from flowers derived from WT Col, WT Ler, *ddm1, kyp2*, *cmt3, ago4 and rdr2* and subjected to PCR to amplify CMT1 RNA sequences using primer set 1+2 (ex10-F+ex11-R) or primer set 3+4 (ex11-F+ex13-R). Note that primer set 3+4 yielded two PCR fragments SP1 and SP2. Actin was used as a reference. PCRs were performed for 35 cycles except for actin (30 cycles). M indicates DNA molecular size markers. The predicted alternative spliced variants SP1 and SP2 are schematically shown below. **c** *CMT1* downstream expression using primer set 5+6 (ex14-F+ex16-R). WT Ler gDNA was used as a reference for PCR product derived from RNA. Note the enhanced downstream *CMT1* expression in *kyp2* and *cmt3* mutants. Actin was used as a reference. Ler genomic DNA (gDNA) was used to confirm amplification from RNA. PCRs were for 35 cycles except for actin (30 cycles). **d** Analysis of splicing out of the entire EK retroelement using primer set 6+7 (ex13-F+ex16-R) on both sides of the EK insertion site. Note lack of a PCR product in *kyp2* and *cmt3* mutant. Actin was used as a reference RNA. Col genomic DNA (gDNA) was used to confirm amplification from RNA. PCR was for 42 cycles except for actin (30 cycles).

Next, we examine *CMT1* expression downstream from the EK insertion site by PCR using primers corresponding to exon 14 (P5) and exon 16 (P6) that expected to yield a PCR product of 227 bp. All lines examined showed recovery of a single PCR product of about the expected size. Yet, the RNA level in WT Col flowers was slightly higher than its level in WT Ler, *ddm1*, *ago4* and *rdr2* mutants. Interestingly, *CMT1* expression downstream from the EK insertion site was significantly enhanced in *cmt3* and *kyp2* mutant flowers (Figure 3C). *CMT1* downstream expression is probably driven by the *CMT1* promoter leading to transcription through the EK retroelement up to the *CMT1* transcription termination site whereby the entire EK can be further spliced out from this chimeric transcript [26]. To test for splicing out of the entire EK transcript, we used a forward primer corresponding to exon 13 (P7) upstream from the EK insertion site and a reverse primer corresponding to exon 16 (P6) downstream from the EK insertion site. If the entire EK has been spliced out it should yield a PCR product of about 268 bp. The results showed (Figure 3D) as expected high *CMT1* expression of this region in WT Col lacking EK. Low level of this region was recovered from WT Ler, *ddm1*, *ago4* and *rdr2* mutants, but no recovery of a PCR fragment could be detected in *cmt3* and *kyp2* mutants (Figure 3D).

Direct sequencing of the resulting PCR products from WT Ler and detailed analysis of the sequencing chromatogram revealed three spliced variants of the entire EK from *CMT1* transcript (Figure 4A); these three spliced variants were confirmed by cloning the PCR products into pJET1.2 and sequencing (Figure 4C-E). The spliced variant SV1 has been described by Henikoff and Comai [26] in which the penultimate base of the EK 3’ LTR serves as a splice donor site, splicing out the entire EK together with a portion of exon 13 and intron 13, to the correct splice acceptor site of exon 14. As a result, the reading frame is shifted resulting in a truncated, non-functional CMT1 protein (Figure 4C). A second spliced variant 2 (SV2) in which the EK is spliced out using a non-canonical donor site (GC) within the duplicated sequence upstream to the EK insertion site (GGCTG-EK) and a canonical AG site contributed by the penultimate base (A) of the EK 5’ LTR and the first base of the duplicated sequence downstream the EK insertion site (EK-GGCTG) resulting in the correct CMT1 reading frame that can potentially produce an intact, functional protein (Figure 4D). The third spliced variant (SV3) in which the entire EK is spliced out using canonical sites (GT-AG) (see Figure 4E) leading to a frame shift and production of a truncated, non-functional CMT1 protein due to a reading frame shift and premature termination (Figure 4E).

**Fig. 4.**
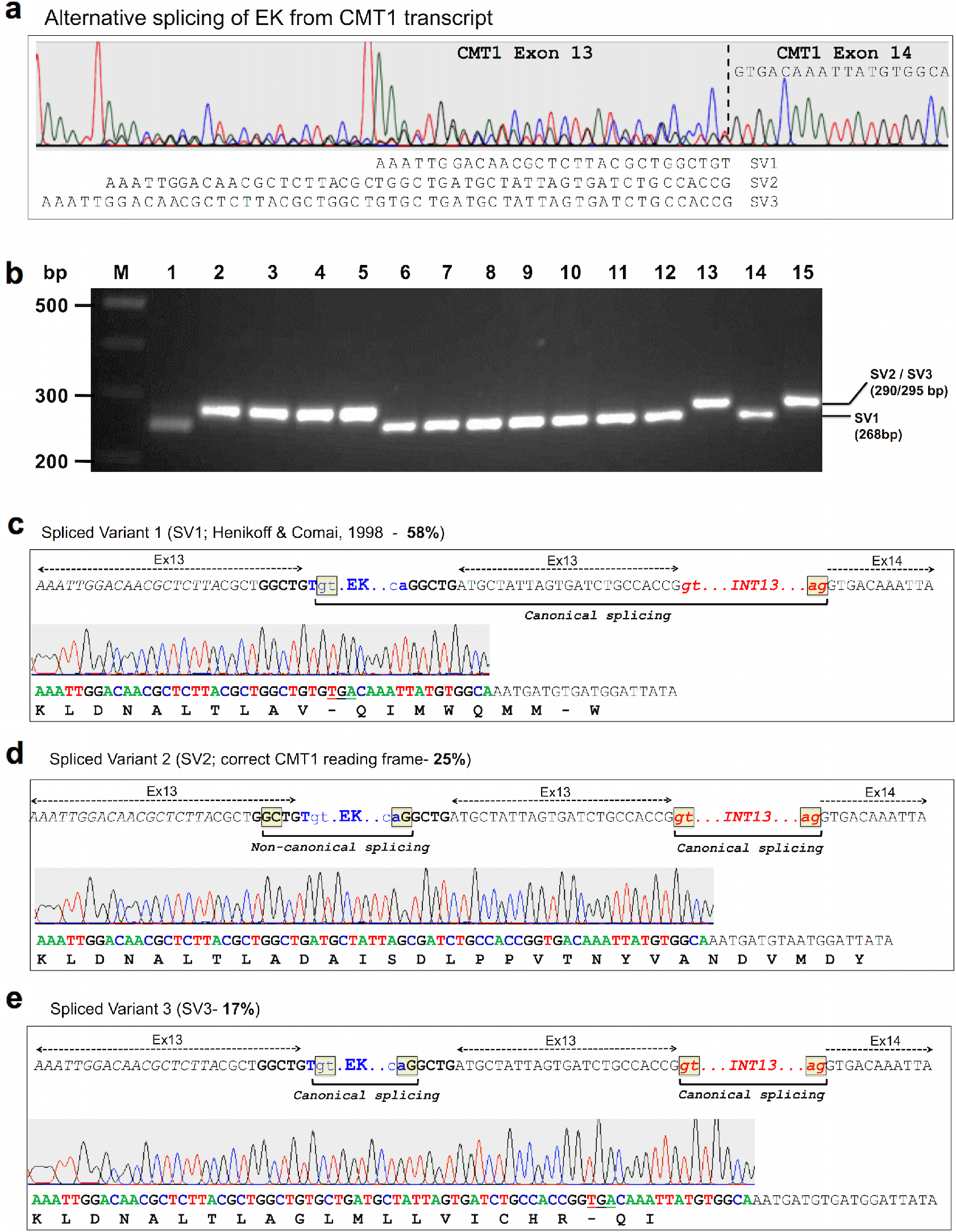
Alternative splicing out of the entire EK in the WT Ler genome. **a** Three alternative variants (SV1, SV2 and SV3) of splicing out of the entire EK revealed by direct sequencing of the PCR product amplified by primers flanking the EK insertion site (P6+P7). Vertical broken line marked the border with exon 14. The sequences of the three spliced variants and of the proximal exon 14 are shown. **b** Analysis of the proportion of spliced variants. An example of PCR analysis of spliced variants (SV1 and SV2+SV3) cloned into pJET1.2 (lanes 1 to 15). Randomly isolated colonies were subjected to PCR followed by separation on 2.5% agarose gel. The position and expected sizes of the various spliced variants are shown on the right. M indicates the DNA size markers. **c** CMT1-EK spliced variant 1 (SV1). The proportion (58%) of SV1 in WT Ler is indicated. The DNA sequence at the EK insertion site is shown with EK marked in blue and intron 13 in red. The fragment spliced out from the chimeric transcript is indicated by a horizontal bracket. The splicing donor (GT) and acceptor (AG) sites are boxed yellow. DNA sequences related to exons 13 and 14 are indicated. The chromatogram and sequence of pJET1.2-SV1 as well as amino acid sequence and premature stop codon (underlined) are shown. **d** CMT1-EK spliced variant 2 (SV2) yielding the correct CMT1 reading frame. The proportion (25%) of SV2 in WT Ler is indicated. The DNA sequence at the EK insertion site is shown with EK marked in blue and intron 13 in red. Horizontal brackets indicate the fragments spliced out from the chimeric transcript. The splicing donor (GC, non-canonical) and acceptor (AG) sites for the EK and the GT-AG canonical splicing sites for intron 13 are boxed yellow. DNA sequences related to exons 13 and 14 are indicated. The chromatogram and sequence of pJET1.2-SV2 and the amino acid sequence are shown. **e** CMT1-EK spliced variant 3 (SV3). The proportion (17%) of SV3 in WT Ler is indicated. The DNA sequence at the EK insertion site is shown with EK marked in blue and intron 13 in red. Horizontal brackets indicate the fragments spliced out from the chimeric transcript. The canonical GT-AG splicing sites are boxed yellow. DNA sequences related to exons 13 and 14 are indicated. The chromatogram and sequence of pJET1.2-SV3 as well as amino acid sequence and premature stop codon (underlined) are shown. The green, red, blue and black peaks in all chromatograms represent the bases ‘A’, ‘T’, ‘C’ and ‘G’, respectively.

To estimate the proportion of the different spliced variants, we cloned the PCR products derived from WT Ler cDNAs into pJET1.2. We randomly isolated colonies and analysed them by PCR followed by separation on agarose gel. Based on the size of the PCR product we could distinguish between SV1 (fast migrating band) and SV2+SV3, the latter variants differ in 5 bp and run on the gel as one band (slow migrating band) (Figure 4B). This analysis revealed 224 positive colonies in which 130 (~58%) were related to SV1 and 94 (~42%) to SV2+SV3. To assess the proportion of SV2 and SV3 variants we sequenced all 94 clones carrying the slow migrating bands (SV2+SV3) and 83 clones produced valid specific sequences. Results showed that 51 (61%) of the clones contain the SV2 fragment and 32 (39%) clones the SV3 fragment. Thus the proportion of the various spliced variants in WT Ler is 58% of SV1, 25% of SV2 and 17% of SV3 (Figure 4C-E).

The possibility that in *cmt3* and *kyp2*, the entire EK is not spliced out from the CMT1-EK-CMT1 transcript was rejected inasmuch as chimeric transcripts having both CMT1 and EK sequences could not be recovered (Supporting information, Figure S3). Thus the enhanced *CMT1* downstream expression in *cmt3* and *kyp2* mutants could have been resulted from EK 5’ LTR functions as a bidirectional promoter, a topic currently studied in the lab.

Finally, we tested for the occurrence of the full length *CMT1* mRNA in WT Ler using RT-PCR. Total RNAs prepared from flowers of WT Col, Wt Ler and *cmt3* (Ler background) were subjected to cDNA synthesis using oligo dT followed by first PCR to amplify *CMT1* coding and 3’UTR regions (Fig. 5A). The full length *CMT1* mRNA appears to be quite rare inasmuch as a faint band of the expected size was visible in Col but not in Ler or *cmt3* mutant (Fig. 5B). However, nested PCR using the first PCR product as template revealed a clear amplified *CMT1* full-length coding region in both Col and Ler wild type ecotypes but not in *cmt3* mutant (Fig. 5B).

**Fig. 5.**
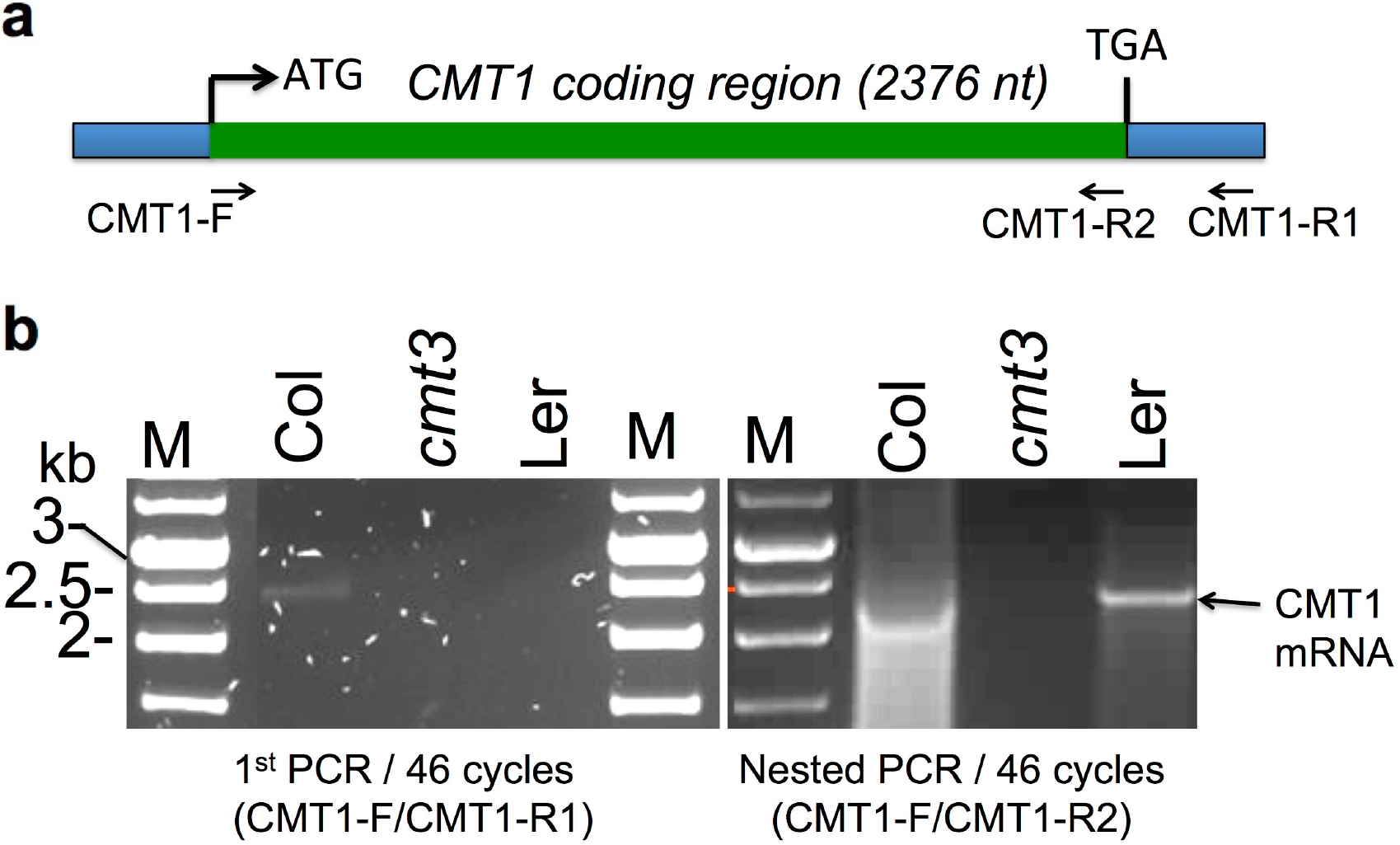
Full length *CMT1* mRNA is produced in WT Ler and Col but not in *cmt3* mutant. **a** Schematic representation of the CMT1 mRNA. The primers used for PCR amplification are shown. **b** RT PCR using cDNAs derived from flowers of WT Col, Wt Ler and *cmt3* (Ler background). The 1^st^ PCR (left panel) was followed by nested PCR (right panel). Arrow indicated CMT1 full length mRNA. M, DNA size markers given in kb.

## Discussion

### Epigenetic control of the exonic Evelknievel retroelement

We showed that the intragenic (exonic) single copy, Evelknievel (EK) retroelement located within exon 13 in the *CMT1* gene is regulated by CMT3 and SUVH4/KYP that cooperate to maintain EK silencing *via* inducing non-CG methylation independently of DDM1 and the RdDM pathway. Thus, in addition to DDM1 and RdDM pathways, our data pointed to CMT3-SUVH4/KYP as an important, independent pathway controlling long TEs located within or near genes at euchromatic regions. Furthermore, silencing of EK by CMT3/KYP is required for splicing out of the entire EK and for reconstitution of a functional *CMT1* mRNA. Thus, our data pointed to an interesting phenomenon whereby the function of CMT1 is rendered partially active by the action of its paralog CMT3 (see model Figure 6). Consequently, retroelement insertion within an exon, does not necessarily lead to complete abolishment of the gene product when the retroelement is kept silent. Rather the retroelement can be spliced out to bring about reconstitution of an intact, functional mRNA. We cannot exclude the possibility that the active CMT1 further participates in methylation and silencing of EK to ensure the persistence of its own expression (see model Figure 6).

**Fig. 6.**
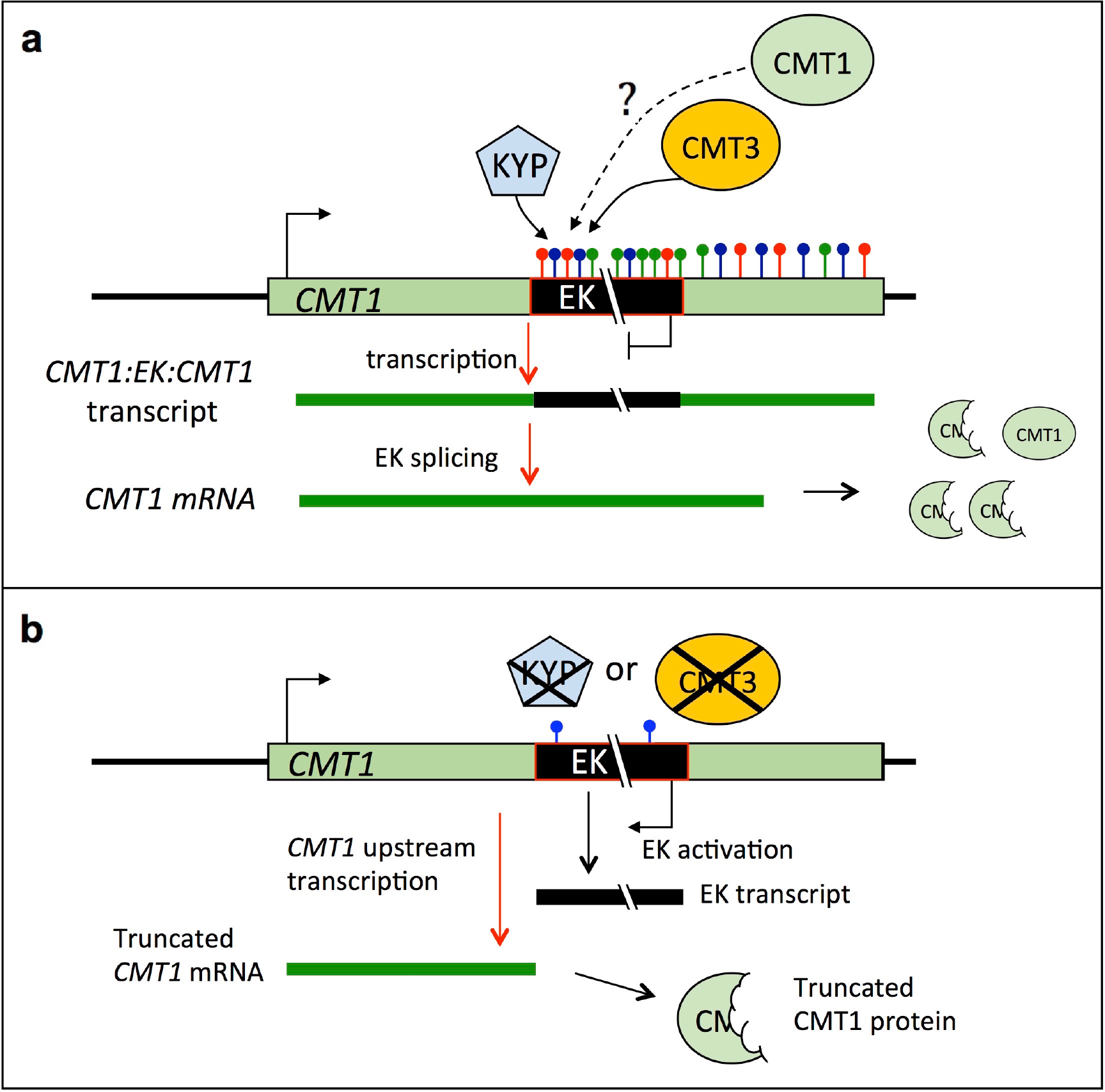
A feed forward model illustrating the regulation of Evelknievel retroelement by CMT3 and CMT1 proteins. **a** Silencing of EK inserted within the *CMT1* gene. CMT3 in concert with SUVH4/KYP maintain cytosine methylation, over the EK retroelement, particularly at the CHG and CHH contexts (red and green lollipops, respectively) and to some extent and in a CMT3-dependent manner also CG methylation (Blue lollipops) as well as histone methylation at lysine 9, which is extended toward *CMT1* 3’ end resulting in a complete silencing of EK. Consequently, transcription from the *CMT1* promoter proceeds through the EK to the *CMT1* transcription termination site. The *CMT1*:EK:*CMT1* transcript undergoes splicing in which EK is spliced out faithfully (in 25% of the events) leading to reformation of mature *CMT1* mRNA and possibly to retrieval of a functional CMT1 protein, which could potentially further methylates EK to ensure its own expression. **b** Activation of EK. In the absence of KYP or CMT3, EK is transcriptionally activated due to absence of CHH and CHG methylation resulting in transcriptional interference of *CMT1* rendering it incapable of transcription beyond the EK insertion site. This leads to formation of a truncated *CMT1* mRNA and to aberrant, non-functional CMT1 protein.

It is commonly accepted that KYP, SUVH5, and SUVH6 H3K9 methyltransferases are required for CMT3-dependent CHG methylation genome wide [10]. Our data clearly showed that redundancy between KYP, SUVH5 and SUVH6 in methylating EK does not exist inasmuch as the sole mutation of KYP was sufficient to drive hypomethylation and expression of EK. Indeed, certain CHG methylation sites are controlled solely by KYP, while mutation in SUVH5 and SUVH6 had no notable effect on CHG methylation [21]. Furthermore, in an early work, Tompa *et al.* [11] performed a genome wide mapping of DNA methylation in *cmt3* mutant *via* fragmentation of the genome with methylation-sensitive enzymes followed by size fractionation and hybridization to microarrays. They identified eight loci displaying reduction in CHG methylation, four of which were found to be low copy number retrotransposons including AtCOPIA10 (At4g21360) and AtCOPIA33 (At2g09830), the later is located at the centromeric region of chromosome 2 near a highly active gene (At2g09990) encoding for 40S ribosomal protein. Thus it appears that the CMT3 and SUVH4/KYP cooperate to maintain non-CG methylation as well as H3K9 methylation and consequently silencing of low copy number, long TEs, which are essentially located near or within active genes. Our analysis of the single copy AtCOPIA18A retroelement (gene ID At5g35935), located on chromosome 5 near active genes (e.g., At5g35970), showed that this element is transcriptionally activated in both *kyp2* and *cmt3* mutants. In a recent article, Sigman and Slotkin [31] proposed that chromosomal location of TEs (i.e., near a gene, within a gene, in a pericentromere/TE island, or at the centromere core) provides the first rule determining the specific regulation of TEs. Accordingly, Sigman and Slotkin [31] proposed that TEs located near genes as well as within genes are initially targeted for DNA methylation by the RNA-directed DNA methylation (RdDM). Once DNA methylation is established, maintenance methylation by MET1 or CMT3 methyltransferases and histone H3 dimethylation by SUVH4/KYP are sufficient to maintain methylation and propagate silencing. Consistent with this proposal is the finding that the EK retroelement inserted within the *CMT1* gene is not activated in RdDM mutants *ago4* and *hen1*, probably because CMT3 and SUVH4/KYP can maintain EK methylation in the absence of RdDM pathway. Similarly, CpG methylation of the 35S promoter sequence was maintained by the activity of MET1 in the absence of RNA trigger [17]. However, unlike CMT3 and SUVH4/KYP, suppression of MET1 activity did not block the establishment of RNA-directed CpG methylation. This raises the question why DNA methylation cannot be re-established at the EK locus by the RdDM pathway in *cmt3* and *kyp2* mutants. One possible explanation is that DNA methylation of the EK retroelement is initially established by the RdDM pathway in conjunction with CMT3 and SUVH4/KYP methyltransferases, rather than DRM methyltransferases. This proposition suggests that CMT3-SUVH4/KYP not only function in maintaining EK non-CpG methylation, which is established by the RdDM pathway, but also in initiating RNA-dependent non-CG methylation at the EK locus. This may gain support by the findings that DRM and CMT3 may act in a complementary manner to maintain RNA-directed non-CG methylation and that mutation of *CMT3* alone could release, to some extent, pre-established RNA-dependent transcriptional silencing of the *NOSpro*:*NPTII* transgene [17]. Alternatively, establishment of non-CG methylation at the EK locus is taking place independently of the RdDM pathway by as yet an unknown mechanism that requires CMT3-SUVH4/KYP pathway. Notably, the RPS (*REPETITIVE PETUNIA SEQUENCE*) from *Petunia hybrida* is efficiently *de novo* methylated, independently of the RdDM pathway, when introduced into heterologous genomes; *RPS* methylation requires the presence of all three DNA methyltransferases MET1, CMT3 and DRM2 [32] and a *cis* regulatory element with the potential of generating stem loop [33].

### Interplay between EK activation and *CMT1* transcription

Commonly TEs are considered as mutable elements in which their insertion into coding or regulatory regions of genes might interfere with proper gene transcription leading to production of aberrant or new transcripts. Yet, plants have evolved various mechanisms to cope with TEs inserted near or within genes [34]. Accordingly, intragenic TEs do not necessarily interfere with proper transcription of the invaded genes due to an IBM2-dependent mechanism that allows synthesis of full length RNA over intragenic TEs carrying repressive epigenetic marks [35, 36]. Our study showed that this is applicable also for exonic TEs. Although EK insertion induces alternative splicing proximal to the insertion site, namely retention of intron 12 (Figure S2), this pattern of *CMT1* upstream transcription is retained whether EK is methylated and silent (WT Ler, *ddm1*, RdDM mutants) or unmethylated and strongly activated (*cmt3* and *kyp2*). Yet, downstream *CMT1* transcription is affected by EK insertion displaying a relatively low level of *CMT1* RNA downstream of the insertion site. Similarly, genes carrying highly methylated intronic TEs displayed defects in transcription downstream of TE insertion site [35]. Methylation and silencing of EK appear to be critical for transcription through the EK element. The chimeric transcript undergoes further processing splicing out the entire EK resulting in a transcript yielding a truncated, non-functional protein [26]. However, our study revealed the existence of two additional spliced variants, namely, SV3 whose splicing follows the canonical GU-AG rule yielding a transcript encoding for a truncated protein, and SV2 whose splicing is non-canonical (GC-AG, [37]) but allows for faithful reconstitution of the *CMT1* RNA that potentially can yield a functional protein. We found that the frequent occurrence of this non-canonical splicing (GC-AG) of the entire EK from *CMT1* transcript is about 25%. Splicing through non-canonical sites (e.g., GC-AG) has been reported both in plants and animals [38,39]. Thus, although it has been proposed previously that CMT1 is dispensable in certain *Arabidopsis* ecotypes carrying EK within the *CMT1* gene (Ler, No-0, RLD, [26]), we cannot exclude the possibility that in these ecotypes a functional *CMT1* mRNA is produced at a very low level (~12.5% considering intron 12 retention) due to splicing out of the entire EK. Indeed, using RT-PCR we could recover low amount of full length *CMT1* mRNA in WT Ler but not in *cmt3* mutant. Whether this level is sufficient for synthesizing a functional CMT1 protein is currently under investigation.

In *cmt3* and *kyp2* mutants *CMT1* downstream transcription driven by the *CMT1* promoter was completely abolished. This can be explained by the oppositely oriented EK and *CMT1* combined with a higher rate of EK transcription resulting in a transcriptional interference [40] that renders *CMT1* incapable of transcription beyond the EK insertion site (Figure 5). Yet, EK activation in *cmt3* and *kyp2* led to strong expression of *CMT1* downstream of the EK insertion site *via* alternative mechanism. We hypothesize that in *cmt3* and *kyp2* mutants the EK 5’ LTR may function as a bidirectional promoter driving both EK and *CMT1* transcription in opposite directions, a topic currently studied in the lab. Precedence for LTRs functioning as bidirectional promoters were described in animals [41-43] as well as in plants [29,44].

### Abbreviations

EK: Evelknievel
CMT: chromomethylase
RdDM: RNA dependent DNA methylation
DRM: domain rearranged methyltransferase
KYP: Kryptonite
MET1: methyltransferase 1
DDM1: decrease in DNA methylation

## Authors’ contributions

NSY designed and performed most of experiments and contributed to results interpretation, preparing figures and writing. JK performed bisulfate sequencing and participated in interpretation and preparation of figures. AZ analyzed methylation data and participated in interpretation and discussion of results. GG conceived of the study, participated in its design and wrote the manuscript. All authors read, commented and approved the final version of the manuscript.

## Acknowledgements

We thank Daphne Autran (IRD, University of Montpllier, France), Caroline Dean (John Innes Centre, UK), Bin Yu (University of Nebraska-Lincoln, USA), Sizolwenkosi Mlotshwa and Vicki Vance (University of South Carolina, USA) and the ABRC for providing mutant lines. A. Cnaani for helping with LTR analysis.

## Ethics approval and consent to participate

Not applicable.

## Supporting information captions

Figure S1: Chop PCR demonstrating erasure of CHG methylation in EK coding region (EKcr) in *cmt3* and *kyp2* mutants

Figure S2: Intron 12 retention

Figure S3: Analysis of chimeric CMT1-EK RNA

Table S1: List of primers used in the present work Bisulfite Sequencing (BS-seq) Data

